# HuR Regulates GATA3-Driven Type 2 Inflammation in CD4^⁺^ T cells and ILC2 in Airway Inflammation

**DOI:** 10.64898/2026.04.23.720195

**Authors:** Fatemeh Fattahi, Laura Yaekle, Julia Holden, Brandon Tepper, Kareem Hussein, Joshua Meier, Liang Xu, Srilaxmi Nerella, Jing Lei, Kelley Bentley, Marc Hershenson, Steven Huang, Ulus Atasoy

**Affiliations:** Division of Allergy and Clinical Immunology, Department of Internal Medicine, University of Michigan Medical School, Ann Arbor, MI; University of Kansas Cancer Center, The University of Kansas Medical Center, Kansas City, KS; Department of Medicine, Division of Pulmonary and Critical Care Medicine, UCSF, San Francisco, CA; Departments of Pediatrics, Molecular and Integrative Physiology, University of Michigan, Ann Arbor, MI; Division of Pulmonary and Critical Care Medicine, Department of Internal Medicine, University of Michigan Medical School, Ann Arbor, MI; Section of Allergy-Immunology, Ann Arbor VA Health System, Ann Arbor, MI

**Author notes:** Corresponding Author: Ulus Atasoy, MD University of Michigan Medical School, Division of Allergy & Clinical Immunology, Department of Internal Medicine 1150 W. Medical Center Dr. MSRB 2, Room 1570B, Ann Arbor, MI 48109-5602.

## Abstract

Type 2–high asthma is driven by coordinated GATA3-dependent programs in CD4⁺ T cells and group 2 innate lymphoid cells (ILC2). Although biologics targeting Th2 cytokines benefit subsets of patients, many remain symptomatic, suggesting upstream regulatory mechanisms may sustain type 2 inflammation. We investigated whether the RNA-binding protein HuR (*ELAVL1*) functions as a post-transcriptional regulator of GATA3-driven type 2 inflammation in allergic asthma. Using a house dust mite (HDM) model *in vivo*, HuR inhibition with KH-3 reduced lung inflammation, suppressed Th2 cytokine expression, accelerated *Gata3* mRNA decay in lung CD4⁺ T cells, and attenuated airway hyperresponsiveness toward control levels. In *ex vivo*–activated human lung CD4⁺ T cells, KH-3 accelerated *GATA3* mRNA decay with minimal effects on *RORC* or *TBX21* and selectively reduced Th2 cytokine secretion, while IL-10 and IL-2 were unchanged. Similarly, ILC2s isolated from PBMCs of type 2–high asthmatic donors showed reduced *GATA3* mRNA stability and diminished Th2 cytokine production following KH-3 treatment. Single-cell transcriptomic analysis of bronchoalveolar lavage fluid after allergen challenge in asthmatic subjects demonstrated co-enrichment of *ELAVL1* and *GATA3* within Th2 clusters in human airways. Together, these findings identify HuR as a therapeutically targetable upstream regulator of GATA3-driven type 2 inflammation in allergic asthma.

## Introduction

Allergic asthma remains a major cause of chronic respiratory morbidity worldwide, affecting approximately 300 million individuals around the world and 25 million in the United States alone, and contributing substantially to the healthcare burden. A substantial proportion of patients exhibit a type 2–high inflammatory endotype, defined by eosinophilic airway inflammation, elevated IgE, and increased production of IL-4, IL-5, and IL-13 (1, 2). These cytokines orchestrate many of the pathologic features of asthma, including eosinophil recruitment, goblet cell metaplasia, mucus hypersecretion, and airway hyperresponsiveness (3). Although inhaled corticosteroids suppress airway inflammation and biologic therapies target IL-4, IL-5, and IL-13 (4, 5), or upstream epithelial alarmins such as TSLP, IL-33, and IL-25 (6–8), a meaningful fraction of patients remains symptomatic, underscoring the need for therapeutic strategies that target upstream or parallel pathways capable of broadly suppressing the inflammatory network driving disease.

Type 2 inflammation is orchestrated by coordinated transcriptional and post-transcriptional programs operating across both adaptive and innate lymphoid compartments (9, 10). GATA3 is the lineage-defining transcription factor for Th2 cells and group 2 innate lymphoid cells (ILC2), driving Th2 differentiation (IL-4/IL-5/IL-13) in CD4⁺ T cells and effector function in ILC2 responding to epithelial alarmins (IL-33/IL-25/TSLP) (11–17). Because GATA3 is a central transcriptional regulator of type 2 inflammatory programs across multiple immune cell types (9, 10), mechanisms that sustain its expression may amplify allergic inflammation; however, the processes that maintain GATA3 abundance under inflammatory conditions remain incompletely defined. Although GATA3 has been extensively studied as a transcriptional regulator (11), far less is known about its post-transcriptional control.

RNA-binding proteins (RBPs) regulate mRNA stability, translation, and localization, thereby shaping both the magnitude and duration of immune responses (18–20). HuR (*ELAVL1*), which binds AU-rich elements (AREs) in the 3′ UTRs of numerous transcripts, is a ubiquitously expressed RBP that stabilizes inflammatory mRNAs and prolongs gene expression (21–24). The *GATA3* mRNA contains AREs within its 3′ UTR, making it a plausible HuR target (25, 26). Consistent with this, our prior work demonstrated that HuR directly binds and stabilizes *GATA3* transcripts in peripheral CD4⁺ T cells from human asthmatic donors, particularly in type 2–high disease, and that T cell–specific HuR ablation in mice significantly attenuates allergic airway inflammation *in vivo*, establishing a causal role for HuR in Th2-driven pathology (26).

Current cytokine-directed therapies act downstream of this broader inflammatory program (4). Because type 2 immunity depends on coordinated expression of lineage-defining and effector transcripts (IL-4/IL-5/IL-13), targeting a shared upstream regulator may provide broader and potentially more durable therapeutic benefit than blocking individual cytokines. KH-3, a small-molecule HuR inhibitor developed by the Xu lab, disrupts HuR–RNA interactions, reduces HuR-mediated transcript stabilization, and has shown *in vivo* activity with acceptable tolerability across diverse murine disease models, including cardiac hypertrophy, nephritis, pancreatic cancer cachexia, and pancreatic cancer metastasis (27–35). These properties support its use as a pharmacologic tool to interrogate HuR-dependent inflammatory pathways *in vivo*.

If HuR is required to maintain GATA3 expression across both adaptive (Th2) and innate (ILC2) type 2 immune compartments (9, 10, 25, 26), pharmacologic inhibition with KH-3 would be expected to shorten GATA3 mRNA half-life, suppress downstream type 2 cytokine production, and attenuate allergic airway inflammation. By targeting a shared post-transcriptional regulator, this approach may overcome a key limitation of existing biologic therapies that act on individual downstream mediators rather than the broader regulatory network.

Here, we tested this hypothesis using complementary murine and human systems, including a house dust mite (HDM)–induced model of allergic airway inflammation, primary murine and human lung CD4⁺ T cells, PBMC-derived ILC2s from asthmatic donors, and single-cell transcriptomic analysis of allergen-challenged human bronchoalveolar lavage fluid. This integrated, multi-level approach allowed us to determine whether the HuR–GATA3 axis is conserved across species and immune compartments and to evaluate whether pharmacologic inhibition of HuR represents a novel upstream therapeutic strategy for type 2–high asthma.

## Results

### HDM-induced type 2 airway inflammation and *Gata3* expression are attenuated by HuR inhibition *in vivo* (Figure 1)

We investigated whether pharmacologic inhibition of HuR modulates established type 2 airway inflammation in an allergen-driven model (**Figure 1A**). At the study endpoint, bronchoalveolar lavage fluid (BALF) and lung tissues were collected for histologic, cytokine, and transcriptional analyses. Histologic evaluation confirmed robust allergic airway inflammation in HDM-challenged mice treated with sham vehicle (HDM+sham), characterized by dense peribronchial and perivascular inflammatory infiltrates, whereas PBS-treated controls showed minimal inflammation (**Figure 1B**). In contrast, KH-3–treated mice showed reduced inflammatory cell infiltration surrounding the airways and blood vessels (**Figure 1B**). Consistent with these findings, HDM challenge induced marked increases in Th2 cytokines. As shown in **Figure 1C**, in BALF, IL-4 and IL-13 were significantly elevated in HDM+sham mice compared with PBS controls and were reduced in HDM+KH-3 mice (*p <* 0.05–0.001 vs. HDM+sham). IL-17 levels, however, were not significantly altered across groups (**Figure 1C**). Similar trends were observed in lung homogenates, where IL-4, IL-5, and IL-13 were significantly increased following HDM exposure and reduced in HDM+KH-3 mice (all \**p <* 0.05 vs. HDM+sham; **Figure 1C**). At the transcriptional level, RT-qPCR analysis of lung tissue showed that HDM exposure increased expression of Th2-associated genes, including *Gata3*, *Il4*, and *Il13* (**Figure 1E**). KH-3 treatment significantly reduced the expression of these transcripts (\**p <* 0.05–0.001 vs. HDM+sham). In contrast, expression of non–Th2 lineage markers (*Rorc*, *Tbx21*, and *Il10*) was not significantly altered, while *Foxp3* showed a modest trend toward increase that did not reach statistical significance (*p =* 0.055) (**Figure 1D**), suggesting selective modulation of the Th2 program by KH-3 treatment. Together, these data demonstrate that pharmacologic inhibition of HuR with KH-3 attenuates HDM-induced type 2 airway inflammation at histologic, molecular, and cellular levels *in vivo*.

**Figure 1.**
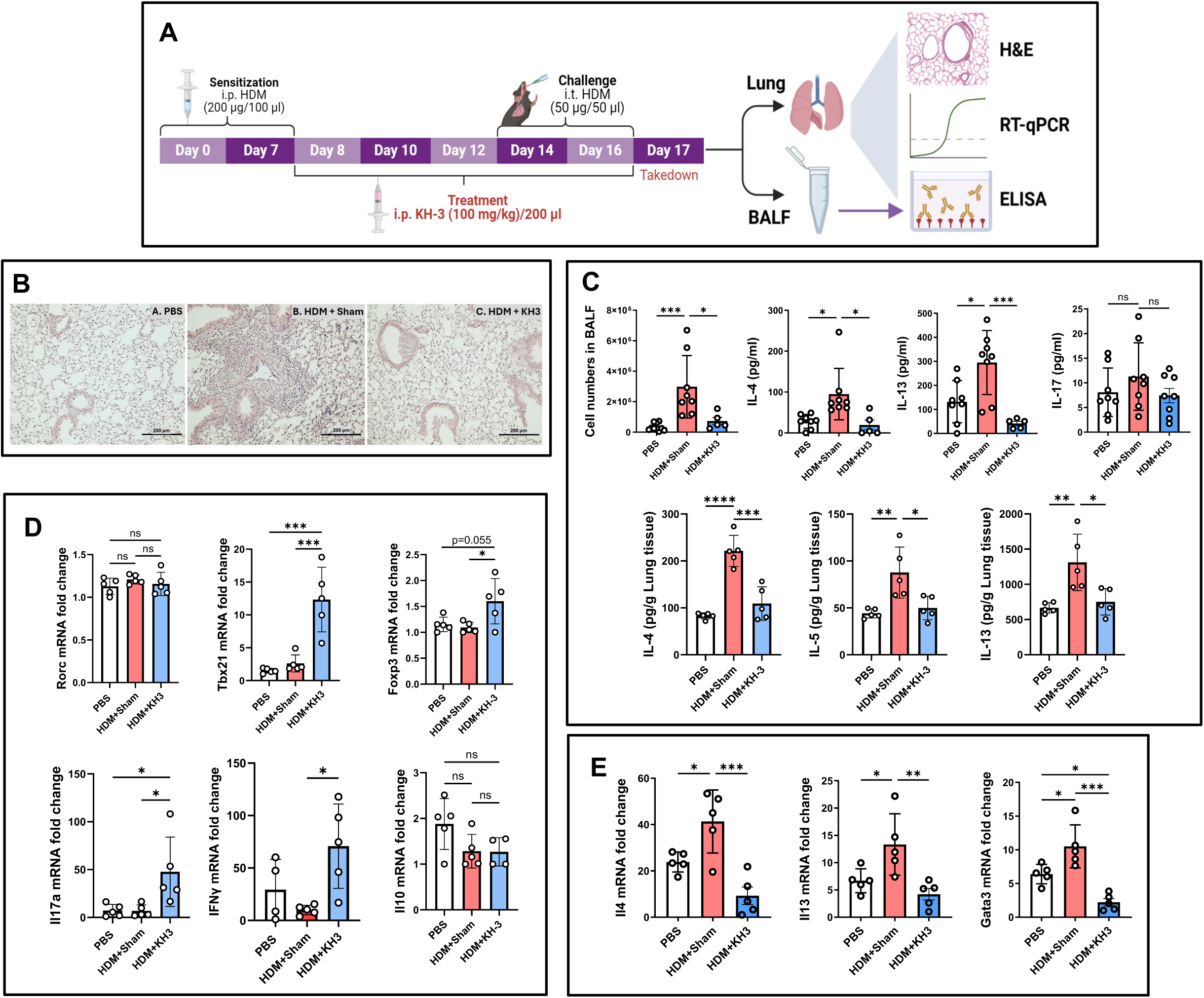
HDM-induced type 2 airway inflammation and *Gata3* expression are attenuated by HuR inhibition *in vivo.* **(A)** Experimental design of the HDM-induced allergic asthma model. Mice were sensitized intraperitoneally with HDM on days 0 and 7 and challenged intratracheally on days 14 and 16. KH-3 was administered intraperitoneally during the challenge phase. Lungs and bronchoalveolar lavage fluid (BALF) were harvested on day 17 for histological, molecular, and cellular analyses. **(B)** Representative hematoxylin and eosin (H&E)–stained lung sections from PBS-, HDM+sham–, and HDM+KH-3–treated mice. HDM+sham mice exhibited marked peribronchial and perivascular inflammatory infiltrates, which were substantially reduced following KH-3 treatment. **(C)** Th2 cytokine protein levels measured by ELISA in BALF (upper panels) and lung homogenates (lower panels). KH-3 significantly reduced IL-4, IL-5, and IL-13 compared with HDM+sham controls. **(D)** RT-qPCR analysis of lung homogenates showing reduced expression of *Gata3* and Th2 cytokines (*Il4*, *Il5*, *Il13*) following KH-3 treatment, with minimal effects on non–Th2 lineage markers. **(E)** BALF cellular analysis demonstrating reduced total leukocyte counts and decreased eosinophils, lymphocytes, and neutrophils in KH-3–treated mice compared with HDM+sham controls. Data are presented as mean ± SD from at least two independent experiments. Statistical analyses in panels (C), (D), and (E) were performed using one-way ANOVA. \**p <* 0.05; \*\**p <* 0.01; \*\*\**p <* 0.001; \*\*\*\**p <* 0.0001.

### HuR inhibition accelerates *Gata3* mRNA decay and diminishes GATA3 protein in allergen-experienced lung CD4⁺ T cells (**Figure 2**)

Because *Gata3* is a central transcriptional driver of Th2 responses, we investigated whether KH-3 suppresses allergic inflammation by destabilizing *Gata3* mRNA in lung CD4⁺ T cells recovered from allergen-challenged mice (**Figure 2A**). As outlined in **Figure 2A**, mice were subjected to HDM sensitization and challenge followed by KH-3 or sham vehicle treatment, after which lung CD4⁺ T cells were isolated and analyzed *ex vivo* by actinomycin D (ActD) chase assays, RT-qPCR, and flow cytometry.

**Figure 2.**
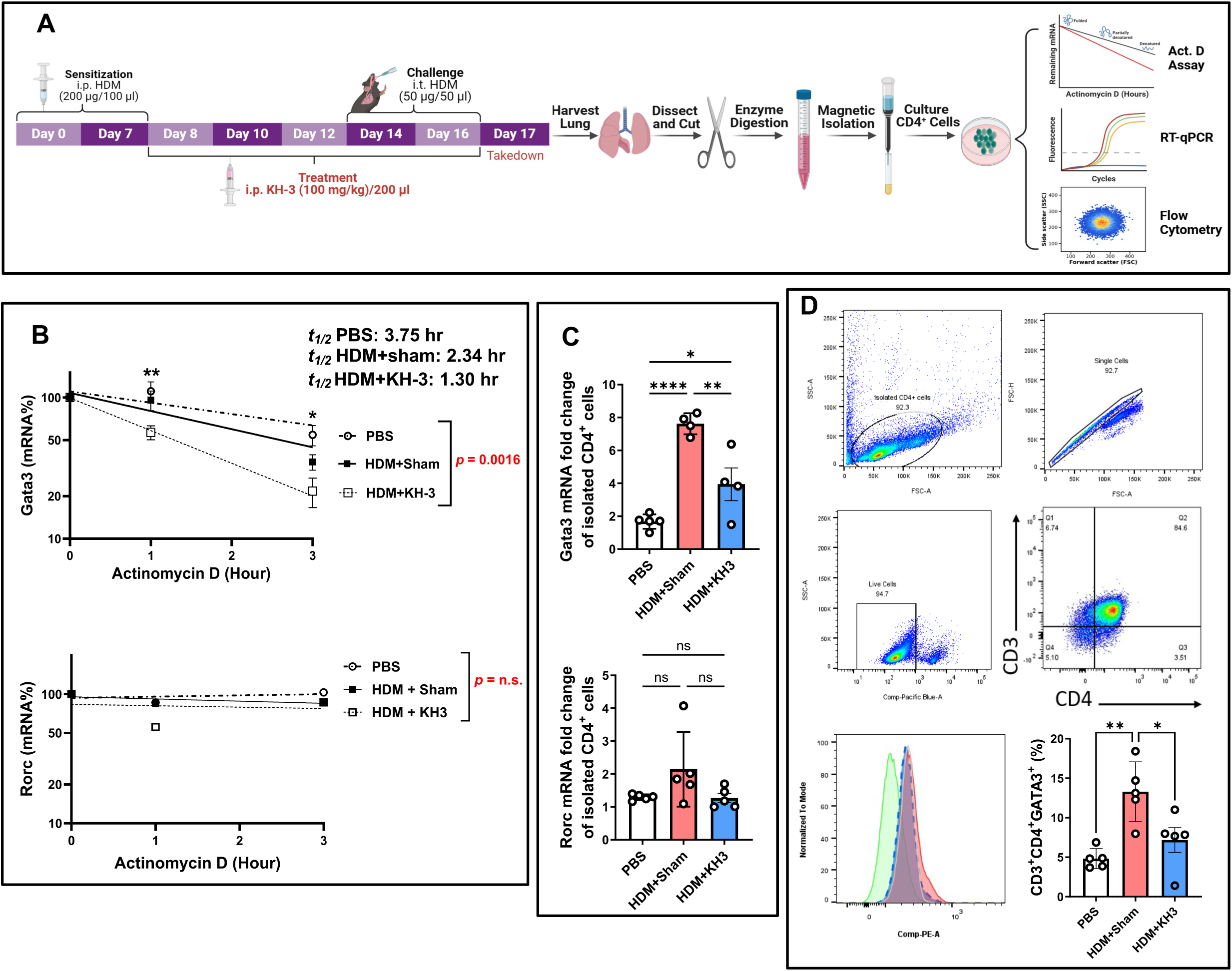
HuR inhibition accelerates *Gata3* mRNA decay and diminishes GATA3 protein in allergen-experienced lung CD4⁺ T cells. **(A)** Experimental schematic. Mice were sensitized and challenged with HDM and treated with KH-3 or vehicle. On day 17, lungs were harvested, enzymatically digested, and CD4⁺ T cells were isolated by magnetic column–based separation. Cells were acutely transferred to culture for actinomycin D (ActD) chase assays, RT-qPCR analysis, or flow cytometric evaluation. **(B)** ActD chase assays demonstrating mRNA stability in isolated lung CD4⁺ T cells. RNA was collected at 1 and 3 hours following ActD treatment. *Gata3* mRNA decay was significantly accelerated in HDM+KH-3–treated mice compared with HDM+sham controls (*t½* values: PBS, 3.75 hr; HDM+sham, 2.34 hr; HDM+KH-3, 1.30 hr; treatment effect *p =* 0.0016). In contrast, *Rorc* mRNA stability was not significantly altered (*p = ns*). Decay curves were analyzed by two-way ANOVA (n = 5 mice per group). **(C)** Steady-state mRNA expression of *Gata3* and *Rorc* in isolated lung CD4⁺ T cells. HDM challenge increased *Gata3* expression, which was significantly reduced by KH-3 treatment, whereas *Rorc* expression remained unchanged. Statistical comparisons were performed using one-way ANOVA (n = 5 per group). **(D)** Representative flow cytometry plots and quantification of CD3⁺CD4⁺GATA3⁺ cells showing reduced GATA3 protein expression in lung CD4⁺ T cells from HDM+KH-3–treated mice compared with HDM+sham controls. Statistical analysis was performed using one-way ANOVA. Data are presented as mean ± SD (n = 5 mice per group). \**p <* 0.05; \*\**p <* 0.01; ns, not significant.

In PBS controls, *Gata3* mRNA decayed gradually following ActD treatment, whereas *Gata3* mRNA turnover was accelerated in the HDM+KH-3 group (**Figure 2B**). Notably, KH-3 treatment further enhanced this decay, resulting in a significantly greater reduction in *Gata3* mRNA over time compared with HDM+sham group (*p =* 0.0016, *t½* values: PBS, 3.75 hr; HDM+sham, 2.34 hr; HDM+KH-3, 1.30 hr). In contrast, *Rorc* mRNA stability was not significantly altered across PBS, HDM+sham, and HDM+KH-3 groups (*p = ns*), indicating that HuR inhibition selectively destabilizes *Gata3* transcripts rather than globally affecting mRNA turnover. Steady-state transcript analysis in purified lung CD4⁺ T cells was consistent with these findings (**Figure 2C**). HDM exposure significantly increased *Gata3* mRNA expression compared with PBS controls, whereas KH-3 treatment reduced *Gata3* transcript levels relative to HDM+sham (*p <* 0.05–0.0001). In contrast, *Rorc* expression remained unchanged across groups (*p = ns*), further supporting selective modulation of the Th2 transcriptional program by KH-3 treatment.

At the protein level, flow cytometric analysis of lung CD3⁺CD4⁺ T cells demonstrated that HDM challenge increased the frequency of GATA3⁺ cells, whereas KH-3 treatment significantly reduced GATA3 expression (**Figure 2D**; *p <* 0.05–0.01 vs. HDM+sham). Representative gating and histograms confirmed a shift toward lower GATA3 expression in KH-3–treated mice.

Together, these data demonstrate that in allergen-experienced lung CD4⁺ T cells, pharmacologic inhibition of HuR with KH-3 accelerates *Gata3* mRNA decay and reduces GATA3 protein expression, supporting a model in which HuR maintains the Th2 transcriptional program by stabilizing *Gata3* transcripts.

### Allergen-induced airway hyperresponsiveness is improved by pharmacologic inhibition of HuR (Figure 3)

To determine whether modulation of HuR, and thereby *Gata3* translates into functional improvement, we assessed airway hyperresponsiveness in the HDM model using invasive plethysmography in intubated, tracheostomized mice (**Figure 3A**). HDM+sham mice exhibited the expected increase in airway responsiveness, with significantly higher airway resistance (R_L_) compared with PBS controls across increasing methacholine doses (**Figure 3B**). KH-3 treatment markedly attenuated this response, resulting in significantly lower airway resistance compared with HDM+sham mice, particularly at higher methacholine concentrations (two-way ANOVA, overall *p <* 0.0001; **Figure 3B**). Airway resistance in KH-3–treated mice were comparable to levels observed in PBS controls. Representative airway resistance tracings of each group demonstrated reduced peak responses in KH-3–treated mice compared with HDM+sham animals (**Figure 3C**). These findings indicate that HuR inhibition by KH-3 improves a key physiological feature of allergic asthma, consistent with its effects on airway inflammation and Th2 responses (**Figures 1** and **2**).

**Figure 3.**
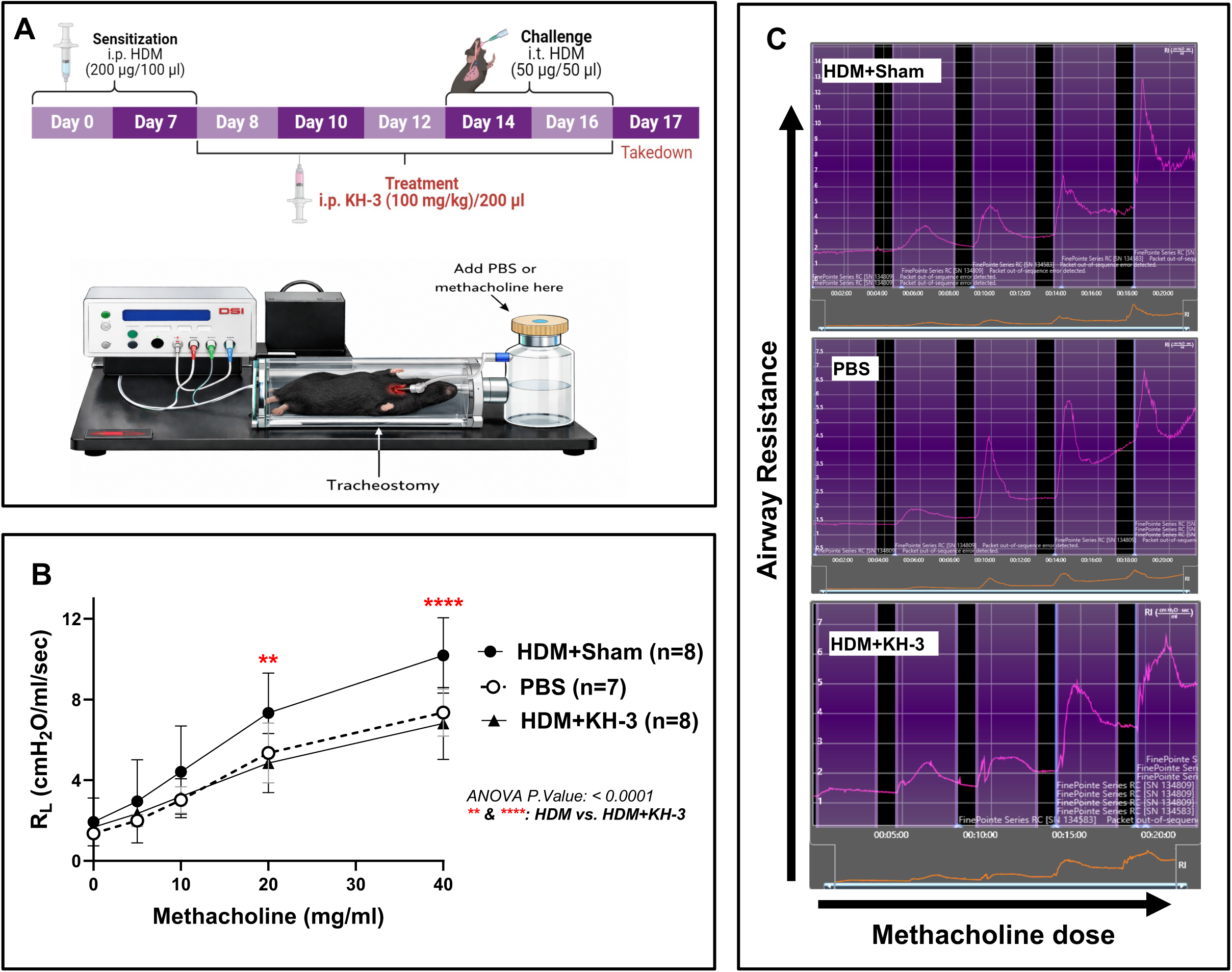
Allergen-induced airway hyperresponsiveness is improved by pharmacologic inhibition of HuR. **(A)** Experimental schematic and setup for measurement of airway resistance using invasive plethysmography following HDM sensitization/challenge and KH-3 treatment. **(B)** Airway resistance (RL) in response to increasing doses of methacholine. HDM+sham mice exhibited exaggerated airway hyperresponsiveness compared with PBS controls, which was significantly attenuated by KH-3 treatment (two-way ANOVA, overall *p <* 0.0001; post hoc comparisons between HDM+sham and HDM+KH-3 are indicated on the graph). Data represent combined results from two independent experiments (PBS, n = 7; HDM+sham, n = 8; HDM+KH-3, n = 8) and are presented as mean ± SD. **(C)** Representative airway resistance tracings from PBS, HDM+sham, and HDM+KH-3 groups across methacholine doses.

### HuR inhibition impairs Th2 effector responses in murine CD4⁺ T cells *in vitro* by (**Figure 4**)

We next asked whether the effects of KH-3 on Th2 programming could be reproduced in a reductionist setting using isolated splenic murine CD4⁺ T cells (**Figure 4A**). As outlined in **Figure 4A**, splenic CD4⁺ T cells were isolated, pretreated with KH-3 or the inactive analog KH-3B, and activated with anti-CD3/CD28, followed by assessment of intracellular cytokine production, transcription factor expression, and cytokine secretion. After 4 days of activation, intracellular staining revealed that KH-3 significantly reduced the proportion of IL-5⁺ and IL-13⁺ CD4⁺ T cells compared with those in KH-3B controls (\**p <* 0.05; **Figure 4B**). IL-17⁺ cells were also reduced (\*\**p <* 0.01), whereas IL-10⁺ and IFN-γ⁺ populations were not significantly altered (*p = ns*). Notably, KH-3 treatment markedly decreased the frequency of GATA3⁺ cells (\*\*\**p <* 0.001), indicating suppression of the Th2 transcriptional program.

**Figure 4.**
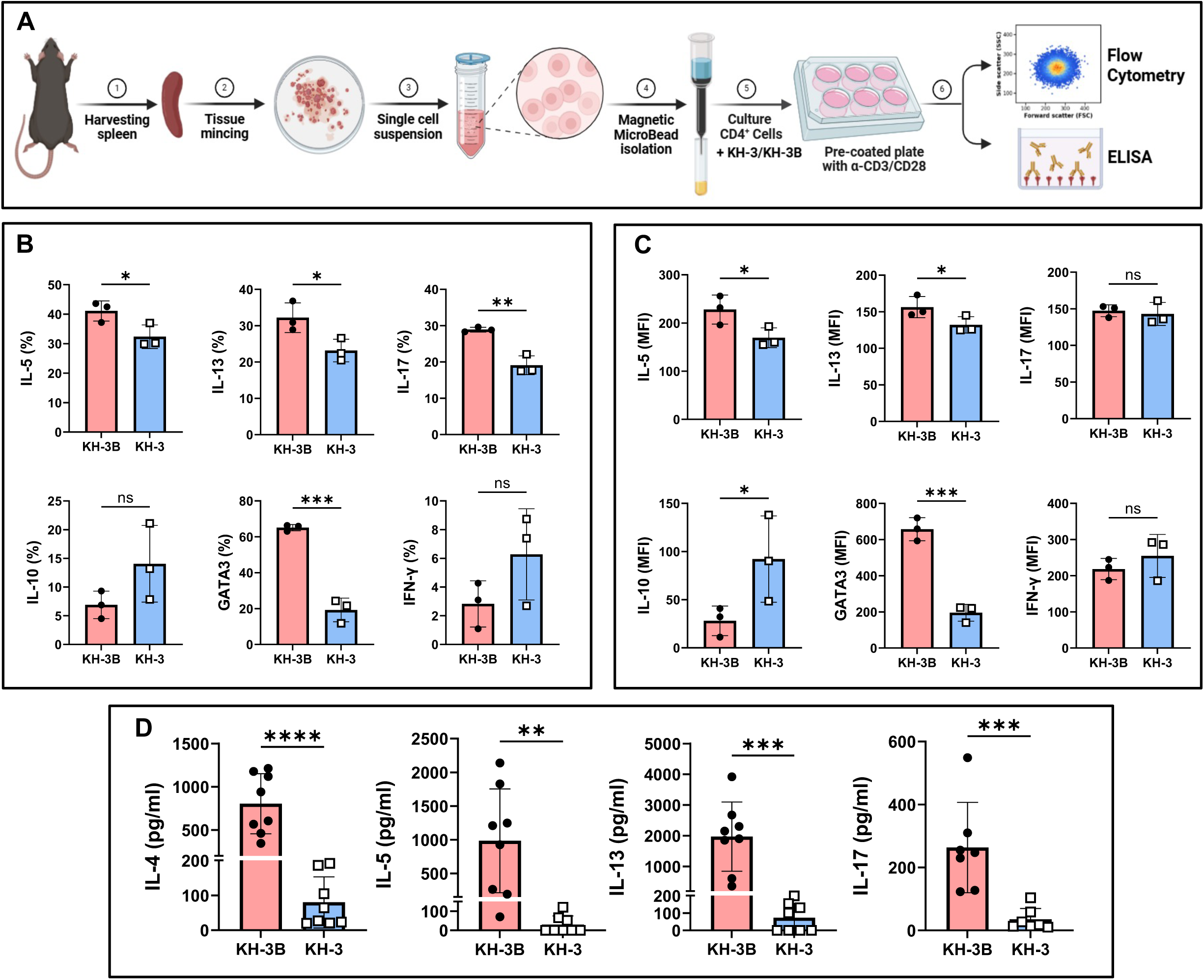
HuR inhibition impairs Th2 effector responses in murine CD4⁺ T cells *in vitro.* **(A)** Experimental workflow. Splenic CD4⁺ T cells were isolated by magnetic bead separation, pretreated with KH-3 or the inactive analogue KH-3B for 2 hours, and then activated with anti-CD3/CD28 for 4 days. Cells were subsequently analyzed by flow cytometry and ELISA. **(B)** Intracellular cytokine staining showing the percentage (%) of IL-5⁺, IL-13⁺, IL-17⁺, IL-10⁺, GATA3⁺, and IFN-γ⁺ CD4⁺ T cells. KH-3 treatment significantly reduced IL-5, IL-13, IL-17, and GATA3 expression compared with KH-3B controls, while IFN-γ and IL-10 were minimally affected. **(C)** Mean fluorescence intensity (MFI) analysis confirming selective suppression of Th2-associated markers following KH-3 treatment. **(D)** ELISA quantification of cytokines in culture supernatants showing reduced secretion of IL-4, IL-5, IL-13, and IL-17 following KH-3 treatment compared with KH-3B controls. Data are presented as mean ± SD from three independent experiments.

Analysis of mean fluorescence intensity (MFI) supported these findings (**Figure 4C**). KH-3 treatment reduced IL-5 and IL-13 expression levels (\**p <* 0.05), and strongly decreased GATA3 MFI (\*\*\**p <* 0.001), while IL-17 and IFN-γ expression remained unchanged (*p = ns*). In contrast, IL-10 MFI was increased following KH-3 treatment (\**p <* 0.05), suggesting a potential shift toward a regulatory phenotype. We next assessed by ELISA the secretion of cytokines that showed significant reduction following KH-3 treatment in the intracellular flow cytometry analyses. Cytokine measurements in culture supernatants were consistent with the intracellular data (**Figure 4D**). KH-3 treatment significantly reduced secretion of IL-4 (\*\*\*\**p <* 0.0001), IL-5 (\*\**p <* 0.01), IL-13 (\*\*\**p <* 0.001), and IL-17 (\*\*\**p <* 0.001) compared with KH-3B controls.

Together, these data demonstrate that pharmacologic inhibition of HuR directly suppresses Th2 effector function in murine CD4⁺ T cells, accompanied by reduced GATA3 expression and cytokine production. These findings are consistent with the *in vivo* data (**Figures 1–3**) and support a model in which HuR acts cell-intrinsically to sustain *Gata3*-dependent Th2 responses.

### HuR inhibition selectively attenuates type 2 cytokine production in *ex vivo*–activated human lung CD4⁺ T cells (**Figure 5**)

To assess whether the HuR–GATA3 axis identified in mouse studies is conserved in human airways, we analyzed CD4⁺ T cells isolated from human lung tissue obtained from healthy donors (**Figure 5A**). As outlined in **Figure 5A**, lung tissue was enzymatically digested to generate single-cell suspensions, followed by magnetic isolation of CD4⁺ T cells. Cells were treated with KH-3 or the inactive analog KH-3B and activated prior to analysis of cytokine production. Using a Luminex platform, we found that KH-3 significantly reduced secretion of key Th2 cytokines, including IL-4 (\*\**p <* 0.01), IL-5 (\*\**p <* 0.01), and IL-13 (\*\*\*\**p <* 0.0001), compared with KH-3B-treated controls (**Figure 5B**). KH-3 also reduced IL-17A (\**p <* 0.05) and IFN-γ (\*\**p <* 0.01), although the extent of suppression was most pronounced for Th2-associated cytokines. However, levels of IL-10, IL-2, and IL-1β were not significantly altered by KH-3 treatment (all *p = ns*), indicating that HuR inhibition does not broadly suppress cytokine production but preferentially targets Th2 responses.

**Figure 5.**
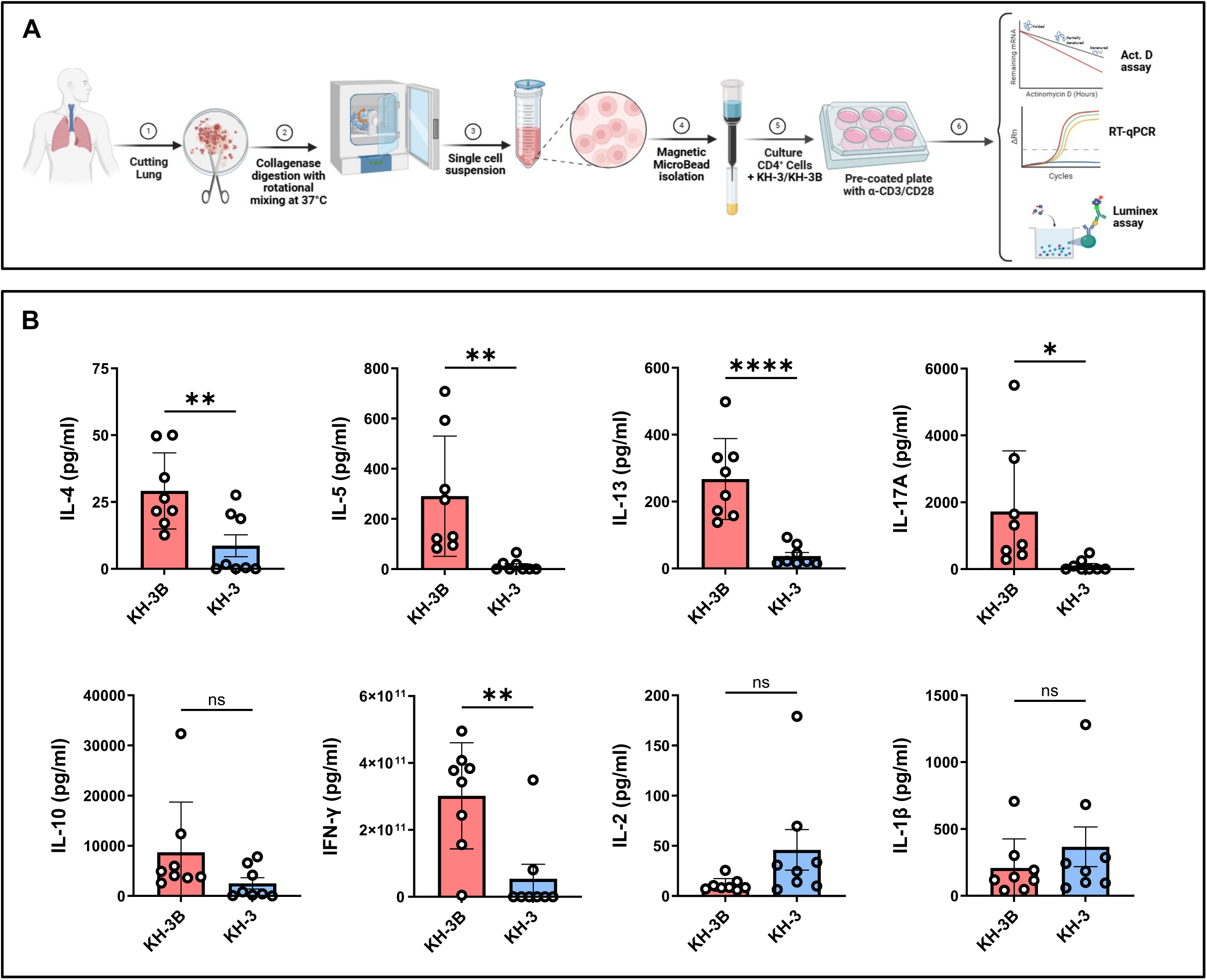
HuR inhibition selectively attenuates type 2 cytokine production in *ex vivo*–activated human lung CD4⁺ T cells. **(A)** Human lung CD4⁺ T cells were isolated from healthy control donors by magnetic bead separation following lung tissue mincing and collagenase digestion. Cells were pretreated with KH-3 or the inactive analogue KH-3B (5 µM) for 2 hours; cells were then activated with anti-CD3/CD28 for 4 days. Culture supernatants were collected and analyzed using a Luminex multiplex assay. Cells were processed for downstream molecular analyses as described in Figure 6. **(B)** KH-3 significantly reduced secretion of IL-4, IL-5, IL-13, IL-17A, and IFN-γ compared with KH-3B controls, whereas IL-10, IL-2, and IL-1β were not significantly changed (as indicated). Data are presented as mean ± SD. Each donor contributed samples from two lung regions (4 total biological samples), and each sample was split into KH-3B vs KH-3 conditions, yielding 8 data points per group. Statistical analyses were performed using paired two-tailed Student’s t test. \**p <* 0.05; \*\**p <* 0.01; \*\*\*\**p <* 0.0001; ns, not significant.

Because each donor contributed paired KH-3B and KH-3 samples, these findings reflect within-donor comparisons and are not driven by inter-individual variability. Together, the data demonstrate that pharmacologic inhibition of HuR selectively attenuates type 2 cytokine production in human lung CD4⁺ T cells, supporting conservation of this pathway across species and implicating GATA3 regulation as a key underlying mechanism.

### HuR blockade destabilizes *GATA3* mRNA and reduces GATA3 expression in *ex vivo*–activated human lung CD4⁺ T cells (**Figure 6**)

To define the mechanism underlying reduced type 2 cytokine production (**Figure 5**), we next examined whether KH-3 affects *GATA3* mRNA stability in human lung CD4⁺ T cells (**Figure 6A**). As shown in **Figure 6A**, CD4⁺ T cells isolated from lung tissue of healthy human donors (n= 4) were pretreated with KH-3 or KH-3B, activated with anti-CD3/CD28, and subjected to ActD chase assays to assess mRNA decay.

**Figure 6.**
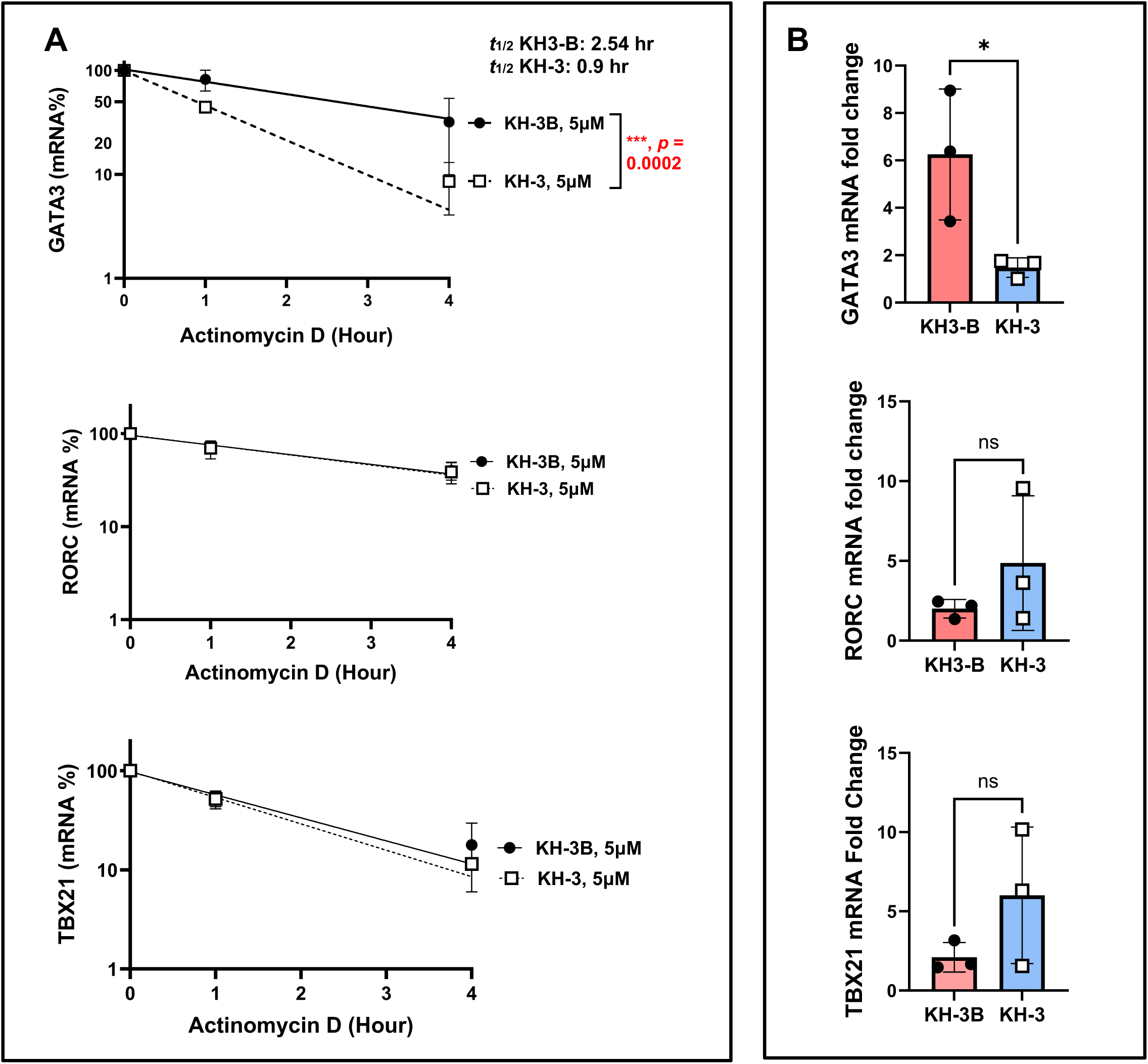
HuR blockade destabilizes *GATA3* mRNA and reduces GATA3 expression in *ex vivo*–activated human lung CD4⁺ T cells. **(A)** Actinomycin D (ActD) chase assays, performed following the experimental design described in Figure 5A, demonstrate accelerated decay of *GATA3* mRNA in KH-3–treated cells compared with KH-3B controls (two-way ANOVA; treatment effect *p =* 0.0002). The calculated mRNA half-life (*t½*) of *GATA3* was 2.54 hours in KH-3B–treated cells and 0.09 hours in KH-3–treated cells, whereas *RORC* and *TBX21* were not significantly altered. **(B)** Steady-state mRNA levels of *GATA3*, *RORC*, and *TBX21* measured by RT-qPCR in activated CD4⁺ T cells after 4 days of stimulation; statistical comparisons were performed using two-tailed Student’s t test. Data are presented as mean ± SD from independent human lung donors. \**p <* 0.05; ns, not significant.

*GATA3* mRNA decayed more rapidly in KH-3–treated cells compared with KH-3B controls, with a significantly greater rate of decline over time. The calculated mRNA half-life (t½) of *GATA3* was 2.54 hours in KH-3B–treated cells and 0.09 hours in KH-3–treated cells (*p =* 0.0002; **Figure 6A**). In contrast, the decay kinetics of *RORC* and *TBX21* were not significantly altered by KH-3 treatment (both *p = ns*), indicating selective destabilization of *GATA3* transcripts.

Consistent with these findings, steady-state RT-qPCR analysis demonstrated that KH-3 significantly reduced *GATA3* mRNA levels in activated human lung CD4⁺ T cells compared with KH-3B controls (\**p <* 0.05; **Figure 6B**). In contrast, expression of *RORC* and *TBX21* remained unchanged (*p = ns*), further supporting selective targeting of the GATA3 transcription factor.

Together, these data demonstrate that pharmacologic inhibition of HuR with KH-3 destabilizes *GATA3* mRNA and reduces its steady-state expression in human lung CD4⁺ T cells, without broadly affecting other lineage-defining transcription factors. These findings mirror the selective effects observed in murine CD4⁺ T cells following HDM challenge (**Figure 2**) and provide a mechanistic basis for reduced type 2 cytokine production in *ex vivo*–activated human airway T cells from healthy donors.

### Type 2 cytokine secretion by human peripheral ILC2s is reduced by HuR inhibition (Figure 7)

Given the growing recognition that ILC2s contribute to persistent and steroid-refractory type 2 inflammation, we next tested whether HuR inhibition affects ILC2 effector responses (**Figure 7**). PBMC-derived ILC2s from healthy controls and type 2–high asthmatic donors were isolated, treated with KH-3 or the inactive analog KH-3B, and stimulated with IL-7, IL-33, and TSLP, as outlined in **Figure 7A**. In cells from asthmatic donors, KH-3 significantly reduced secretion of TNF-α (\*\**p <* 0.01) and IL-13 (\**p <* 0.05) (**Figure 7B**). IL-4, IL-6, and GM-CSF showed a trend toward reduction, with p values indicated in **Figure 7B**, whereas IL-5, IL-9, MIP-3α, and IL-15 were not significantly altered (*p = ns*). Paired analyses within each donor demonstrated a consistent downward shift in canonical type 2 cytokines with KH-3 treatment, whereas non–type 2 mediators were relatively preserved. These findings indicate that HuR supports type 2 effector programs in human ILC2s, consistent with a similar pattern observed in human lung Th2 cells (**Figure 5**).

**Figure 7.**
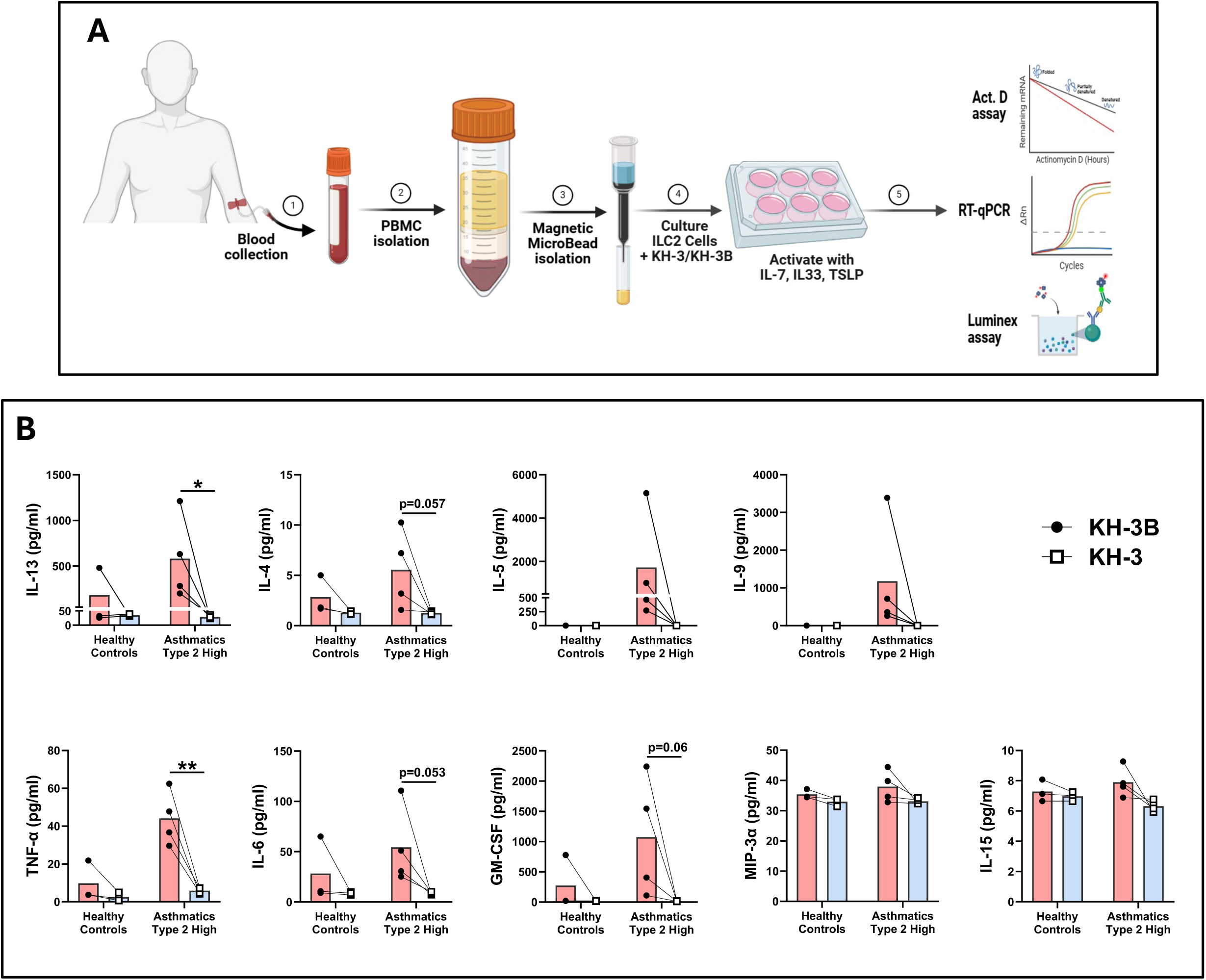
Type 2 cytokine secretion by human peripheral ILC2s is reduced by HuR inhibition. **(A)** Human PBMC-derived ILC2 were isolated and treated with KH-3 or the inactive analogue KH-3B, then activated with IL-7, IL-33, and TSLP prior to supernatant collection for Luminex analysis and cells were analyzed for molecular assay as will described in Figure 8. **(B)** KH-3 reduced cytokine production in type 2–high asthmatic donors, including IL-13, IL-4, IL-5, IL-9, TNF-α, IL-6, and GM-CSF, with minimal effects on MIP-3α and IL-15. Responses are shown for healthy controls (n = 3) and type 2–high asthmatic (n = 4) donors, with paired lines indicating matched conditions within each donor. Statistical comparisons were performed using paired two-tailed Student’s t test; exact p values are shown on the graphs where indicated. Data are presented as mean ± SD. \**p <* 0.05; \*\**p <* 0.01; ns, not significant.

### Decreased *GATA3* mRNA half-life and altered ILC2 effector-associated transcripts following HuR inhibition (Figure 8)

To determine whether HuR regulates GATA3 post-transcriptionally in human ILC2s, we performed ActD chase assays on PBMC-derived ILC2s from type 2–high asthmatic donors treated with KH-3 or KH-3B (**Figure 8A**). *GATA3* mRNA decayed more rapidly in KH-3–treated ILC2s than in KH-3B controls, with the estimated half-life decreasing from 1.19 hours under KH-3B conditions to 0.66 hours with KH-3 (*p <* 0.0001, two-way ANOVA; **Figure 8A**), paralleling the KH-3–induced destabilization of GATA3 observed in human lung CD4⁺ T cells (**Figure 6B**).

**Figure 8.**
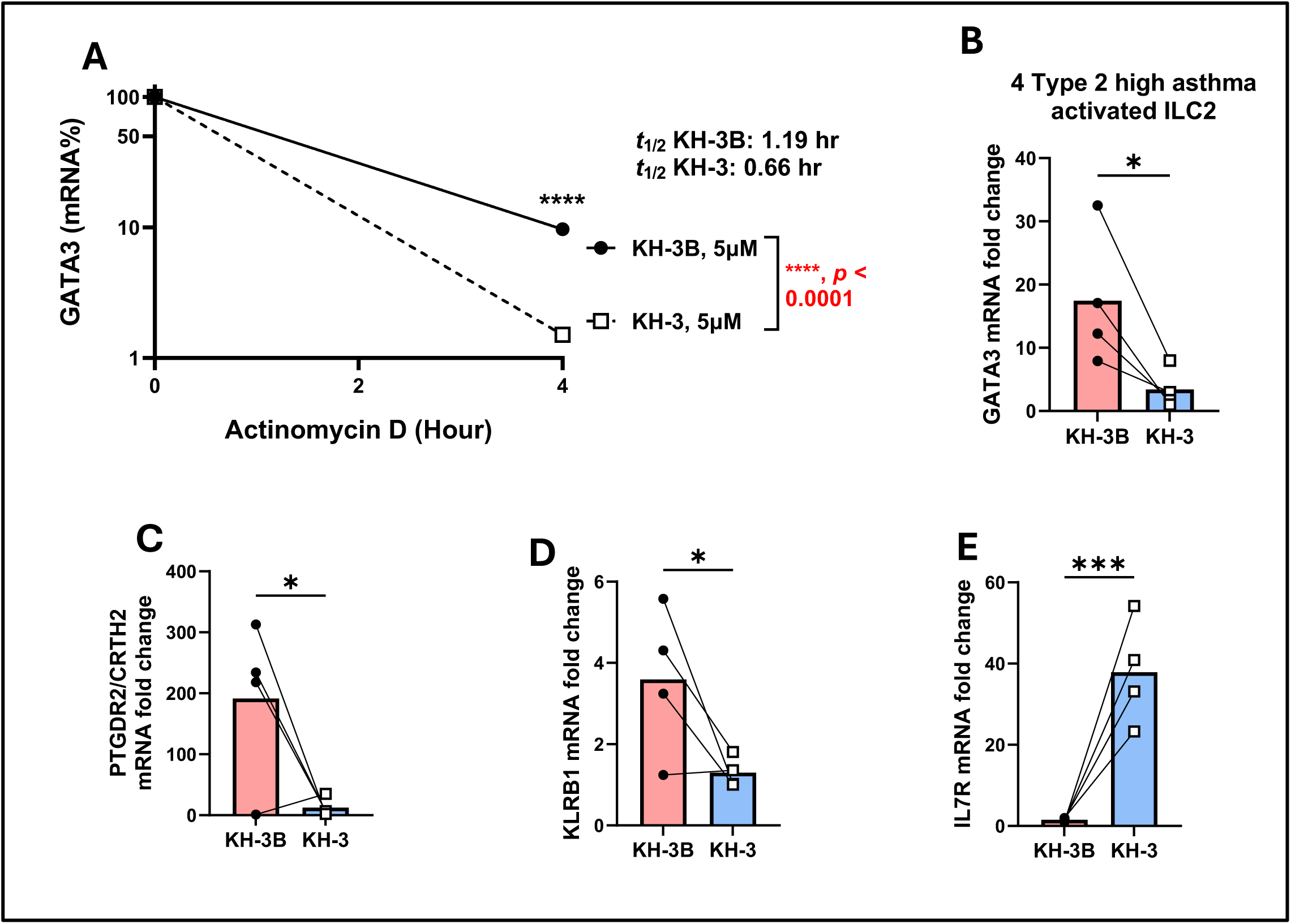
Decreased *GATA3* mRNA half-life and altered ILC2 effector–associated transcripts following HuR inhibition. Actinomycin D (ActD) chase assays, performed following the experimental design described in Figure 7A, demonstrate accelerated decay of *GATA3* mRNA following KH-3 treatment (t½: KH-3B, 1.19 hr; KH-3, 0.66 hr; \*\*\*\**p <* 0.0001, two-way ANOVA). **(B–E)** Steady-state mRNA expression of *GATA3*, *PTGDR2*, *KLRB1*, and *IL17R* measured by RT-qPCR. KH-3 significantly reduced *GATA3*, *PTGDR2*, and *KLRB1*, while *IL17R* expression was increased. Data are presented as mean ± SD from independent donors. Statistical comparisons were performed using paired two-tailed Student’s t test. \**p <* 0.05; \*\*\**p <* 0.001; ns, not significant.

We next assessed steady-state expression of ILC2-associated genes after HuR inhibition. KH-3 significantly reduced *GATA3* mRNA abundance compared with KH-3B (*p <* 0.05; **Figure 8B**) and lowered expression of *PTGDR2* (encoding CRTH2) and *KLRB1* (encoding CD161) (both *p <* 0.05; **Figure 8C, 8D**), consistent with a loss of the canonical type 2 ILC2 phenotype. In contrast, *IL7R* (encoding the IL-7 receptor α chain, CD127) showed a significant increase in steady-state mRNA after KH-3 treatment (*p <* 0.001; **Figure 8E**), indicating that not all ILC2-associated transcripts are uniformly suppressed by HuR inhibition. Together with the Luminex data (**Figure 7**), these findings indicate that KH-3 constrains ILC2 effector function primarily through destabilization of *GATA3* mRNA and suppression of canonical type 2 ILC2-associated programs.

### Single-cell transcriptomic analysis of BALF links *ELAVL1* and *GATA3* expression to Th2 cells in human allergic airways following allergen challenge (Figure 9)

To place these mechanistic observations in the context of *in vivo* human airway inflammation, we analyzed single-cell RNA sequencing (scRNA-seq) data from BALF obtained after segmental HDM or control challenge (**Figure 9A**). Unsupervised clustering and UMAP visualization resolved major immune lineages, including CD4⁺ and CD8⁺ T cell subsets, B cells, ILC2, myeloid cells, mast cells, and other cell populations (**Figure 9B**).

**Figure 9.**
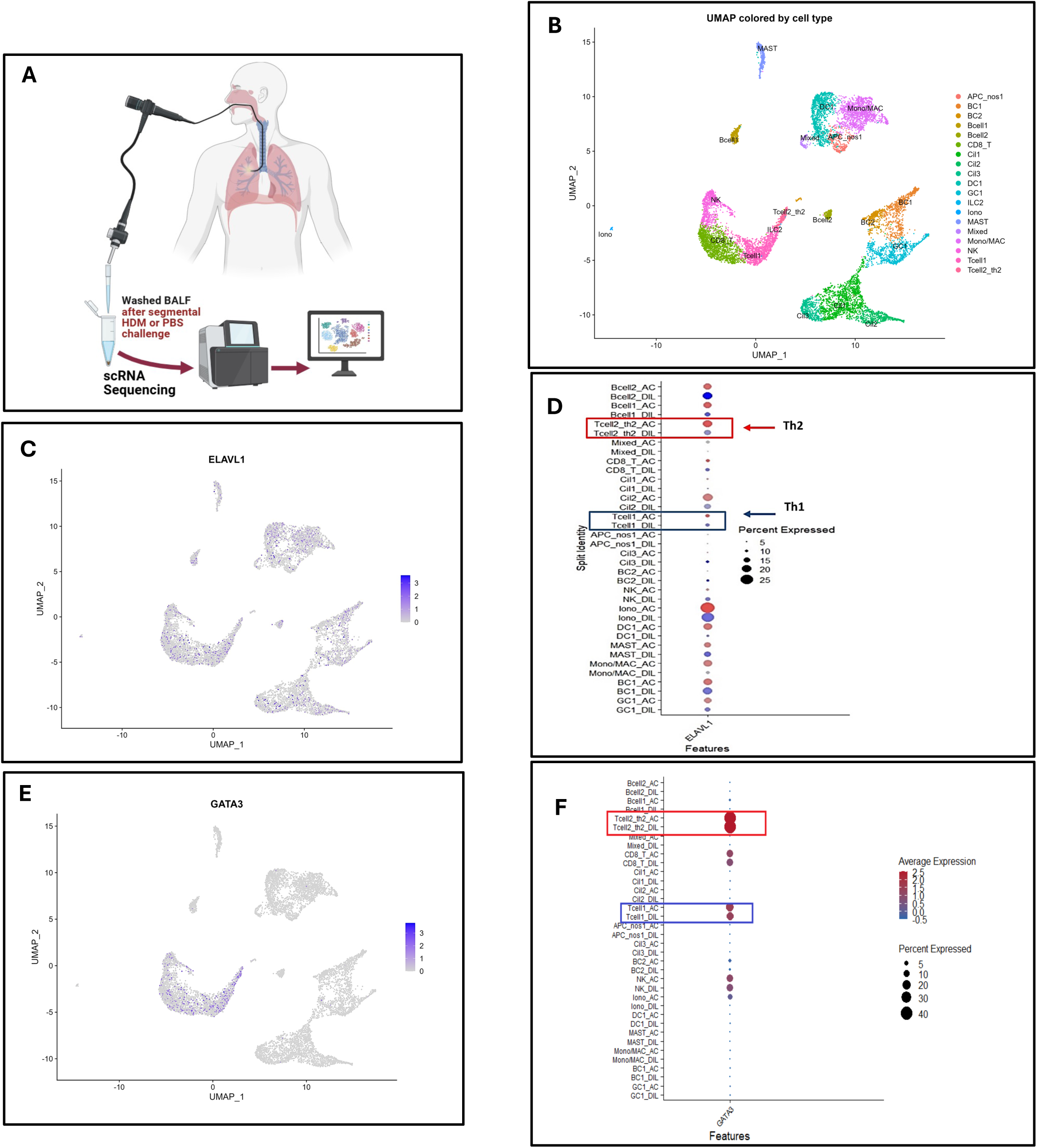
Single-cell transcriptomic analysis of BALF links *ELAVL1* and *GATA3* expression to Th2 cells in human allergic airways following allergen challenge. **(A)** Schematic of experimental workflow. Bronchoalveolar lavage fluid (BALF) was collected following segmental HDM or PBS challenge. Washed BALF cells were processed for single-cell RNA sequencing (scRNA-seq) to characterize airway immune cell populations. **(B)** UMAP visualization of scRNA-seq data colored by annotated immune cell populations, including CD4⁺ T cell subsets, CD8⁺ T cells, B cells, ILC2, dendritic cells, monocytes/macrophages, mast cells, and other immune clusters. Distinct clustering confirms successful resolution of major airway immune populations. **(C)** Feature plot showing expression of *ELAVL1* (encoding HuR) across immune cell populations, demonstrating broad expression with enrichment in T cell clusters. **(D)** Dot plot summarizing *ELAVL1* expression across annotated cell types in HDM-challenged and diluent control samples, showing increased expression in Th2-like CD4⁺ T cell clusters relative to other lymphocyte subsets. **(E)** Feature plot displaying *GATA3* expression across the UMAP projection, localized predominantly to Th2-like CD4⁺ T cell populations. **(F)** Dot plot summarizing *GATA3* expression across immune cell clusters, demonstrating enrichment in Th2 cells and limited expression in non–type 2 immune subsets. Data represent integrated scRNA-seq analysis of BALF immune cells from HDM- and PBS-challenged human airways. Dot size reflects the percentage of cells expressing the gene, and color intensity represents average expression.

*ELAVL1*, the gene encoding HuR, was broadly expressed across immune cells but showed higher expression in T cell–enriched clusters (**Figure 9C**). Dot plot analysis further confirmed enrichment of *ELAVL1* in Th2-like CD4⁺ T cell clusters relative to other lymphocyte subsets (**Figure 9D**). Feature and dot plot analyses of GATA3 demonstrated that its expression was largely confined to Th2-like CD4⁺ T cells (**Figure 9E, F**). The overlap of *ELAVL1* and *GATA3* in these Th2-skewed clusters is consistent with our functional data showing that HuR stabilizes *GATA3* mRNA and supports Th2 effector function in murine and human CD4⁺ T cells (**Figures 2, 4**, and **5**). While the single-cell data in Figure 9 are correlative and do not establish causality, they provide in situ evidence that *ELAVL1* and *GATA3* are co-expressed in CD4⁺ T cell populations that contribute to allergic airway inflammation.

## Discussion

Type 2–high asthma arises from coordinated activation of adaptive Th2 cells and innate ILC2s, yet current biologic therapies predominantly neutralize individual effector cytokines or their receptors rather than upstream mechanisms that restrain both arms of type 2 immunity, and many patients remain symptomatic despite treatment (5–8, 36–38). Here, we identify the RNA-binding protein HuR as an upstream post-transcriptional regulator of this shared type 2 program and show that inhibiting HuR reduces *GATA3* mRNA stability, dampens Th2 and ILC2 cytokine production, and improves allergen-induced airway disease in complementary mouse and human systems (25, 26, 39). These findings support post-transcriptional control of GATA3-dependent circuits as an additional layer of regulation in allergic asthma beyond current cytokine-targeted biologics (6, 26, 36, 40).

Prior studies, including our own, demonstrated that HuR stabilizes *GATA3* transcripts and promotes Th2 cytokine production, mainly using T cell–specific HuR deletion or siRNA in peripheral blood–derived T cells (25, 26, 40, 41). HuR overexpression enhances GATA3 expression and Th2 cytokine production, whereas HuR silencing reduces *GATA3* mRNA stability and attenuates downstream cytokine expression, supporting a central role for HuR in coordinating the Th2 program (25). Here we show that a drug-like small molecule can similarly destabilize murine *Gata3* and human *GATA3* mRNA and suppress downstream Th2 cytokines in both murine and human lung CD4⁺ T cells. This supports a model in which GATA3-driven responses are sustained not only by lineage commitment but also by continued RNA stabilization, with HuR acting as a molecular scaffold that maintains the type 2 transcriptional program in inflamed tissue (19, 22, 25, 26). *In vivo*, KH-3 reduced BALF inflammatory cellularity, airway inflammation, and lung Th2 cytokines, and it improved airway hyperresponsiveness in a HDM-driven model, consistent with prior HuR-loss studies in the T-cell and epithelial compartments of allergen-exposed mice (25, 26, 39–41).

Our human studies demonstrate HuR-dependent regulation of type 2 pathways in primary lung CD4⁺ T cells and circulating ILC2s from asthmatic donors, providing mechanistic insight in disease-relevant lymphocyte populations (26, 38). As direct access to human airway tissue is limited (42), combining lung-derived CD4⁺ T cells with PBMC-derived ILC2s provides a practical window into HuR-regulated type 2 immunity in humans.

Segmental allergen challenge studies have demonstrated dynamic airway responses to allergen exposure and enrichment of type 2 effector populations in BALF from asthmatic patients (43–47). To place our HuR findings in this context, we reanalyzed the single-cell BALF dataset generated by Siddiqui *et al.* (47), which revealed enrichment of *GATA3*- and *ELAVL1*-associated transcripts in Th2-skewed CD4⁺ T-cell clusters, suggesting activation of type 2–related regulatory programs in the human airway compartment (26, 46). Together with our finding that KH-3 accelerates *GATA3* mRNA decay and reduces Th2 cytokine production in lung-derived CD4⁺ T cells from asthmatic donors, these observations provide evidence that the HuR–GATA3 axis operates in the relevant anatomic compartment in human disease (26).

Consistent with classic studies showing that CD4⁺, but not CD8⁺, T cells are the dominant source of Th2-type cytokine mRNA and eosinophil-prolonging factors in blood and BALF of asthmatic patients, we observed broad *ELAVL1* expression with prominent enrichment in GATA3⁺ CD4⁺ T-cell clusters (43, 44, 48). Although these data remain associative, they provide a strong rationale for future perturbation studies combining HuR inhibition with single-cell multiomics to define the HuR-dependent transcriptional network in human airway CD4⁺ T cells, building on existing single-cell maps of allergen-challenged airways (46). These observations further suggest that HuR targeting could complement current biologics and optimized multi-drug inhaler regimens, particularly in patients whose disease remains active despite IL-5, IL-4Rα, or TSLP blockade and single- or triple-inhaler controller therapy (6, 7, 36).

Our results also broaden HuR biology beyond adaptive T cells by implicating it in ILC2 function (26, 39, 49). ILC2s are now recognized as central drivers of type 2 inflammation, especially in exacerbations and disease recurrence, and they cooperate with CD4⁺ T cells to amplify airway pathology (8, 37, 38, 50, 51). Several groups have highlighted their distinct transcriptional and metabolic dependencies, including GATA3- and c-Myc–centered programs and IL-33–driven developmental checkpoints (9–11, 52–57). In particular, an ILC2-restricted tandem super-enhancer is required for stage-specific GATA3 up-regulation and ILC2 maintenance, while sparing Th2 cells (52, 53). We found that KH-3 reduced IL-13, IL-4, and additional inflammatory mediators in PBMC-derived ILC2s from asthmatic donors, indicating that HuR supports a shared effector program across Th2 cells and ILC2s rather than a T-cell-restricted pathway. Given accumulating evidence that targeting ILC2s, either directly or via upstream mediators such as TSLP, IL-33, or CRTh2, can mitigate asthma relapse and steroid-refractory disease (8, 37, 50–52, 54, 55, 58, 59), HuR inhibition may offer a way to simultaneously restrain both adaptive and innate type 2 compartments.

An additional strength of HuR inhibition in our system is its preferential effect on type 2–associated programs. Across our models, HuR blockade using KH-3 had modest effects on non–type 2 markers such as *TBX21*, *RORC*, IL-2, or IL-10, consistent with preferential regulation of AU-rich 3′UTR–containing transcripts rather than uniform effects across the transcriptome (19, 21, 22, 25, 26, 49). Similar selectivity has been reported in other disease models, where KH-3 or related HuR inhibitors attenuate pathogenic gene expression in cancer, cardiac remodeling, and kidney disease without broad side effects or cytotoxicity (27, 30–33, 60). For chronic diseases like asthma, where long-term safety is critical and many patients already receive triple inhaler therapy, the ability to blunt type 2 pathways without broadly paralyzing immune function will be essential (4, 6, 36). Our findings also link pharmacologic HuR inhibition to our prior genetic studies (26, 41). T cell–specific HuR ablation impairs Th2 differentiation and reduces allergen-driven lung inflammation (26, 41), whereas HuR deficiency in regulatory T cells destabilizes *Foxp3* and compromises suppressive function (49). In the present study, KH-3 recapitulated key features of the Th2-directed HuR knockout, including reduced type 2 cytokine production and improved airway physiology, without directly affecting *Foxp3* expression in murine lung in our experimental systems (26). Together, these observations indicate that HuR regulates CD4⁺ T-cell responses in a context-dependent manner, with a preferential effect on type 2 effector responses in allergic airway inflammation models. Along with reports that HuR controls chemokine production in airway epithelial and smooth muscle cells and contributes to severe asthma-like phenotypes (39), our findings underscore the broad influence of HuR across both structural and immune compartments in the lung and support further evaluation of airway-targeted HuR modulation as a potential strategy to dampen pathogenic type 2 responses while carefully monitoring effects on regulatory pathways (26, 39).

Several limitations should be considered when interpreting our findings. Although KH-3 demonstrated consistent activity across multiple murine models and *ex vivo* human lymphocyte systems, additional studies will be important to further define its pharmacologic properties, tissue distribution, and long-term safety profile *in vivo* (27, 30–32, 60). Moreover, while our mechanistic studies focused on *GATA3* as a central HuR-regulated target, HuR is known to interact with numerous inflammatory transcripts, and coordinated regulation of multiple pathways likely contributes to the overall phenotype observed after HuR inhibition (19–22, 25, 26, 39, 61). Future transcriptomic and translational profiling studies, including polysome- and ribosome-profiling approaches, will be important to further define the broader HuR-dependent regulatory network in type 2 airway inflammation. Our human cohorts were modest in size and enriched for donors undergoing clinically indicated procedures, which may limit generalizability (42, 46). In addition, the BALF single-cell analyses are associative and do not directly establish a causal requirement for HuR activity in maintaining airway Th2 and ILC2 populations *in vivo*.

Future studies integrating HuR inhibition with single-cell multiomic or perturbational approaches may help further define cell-type-specific regulatory programs and therapeutic vulnerabilities (7, 46, 56, 60). Despite these limitations, the present study provides evidence across complementary mouse and human systems that pharmacologic HuR inhibition destabilizes GATA3-associated type 2 programs and attenuates allergic airway inflammation. These findings support continued evaluation of HuR-targeted strategies, including airway-directed delivery approaches and studies in steroid-refractory or exacerbation-prone asthma settings, where type 2 effector populations remain important drivers of disease (6, 36, 46, 51). It will also be important to further define how HuR intersects with additional stress-response and metabolic pathways that influence Th2 and ILC2 survival, activation, and plasticity, including pathways linked to RBM3, ferroptosis, and mitochondrial fitness (6, 36, 51, 52, 56, 57, 62).

In summary, our work adds to a growing body of evidence that RNA-binding proteins are critical, yet still underexplored, regulators of immune pathology (19, 20, 56). By destabilizing *GATA3* and additional type 2-associated transcripts, HuR inhibition with KH-3 attenuates Th2 and ILC2 effector function and improves allergen-driven airway disease *in vivo* (25, 26, 39, 40). These findings suggest that targeting post-transcriptional regulatory mechanisms could complement current cytokine-blocking biologics and advanced inhaled therapies, offering a broader strategy for the treatment of type 2-high asthma and related airway inflammatory diseases.

## Materials and Methods

### Study design and randomization/blinding

This study was designed to determine whether the RNA-binding protein HuR regulates GATA3-dependent type 2 inflammatory responses characteristic of allergic asthma. To interrogate HuR function, we used the small-molecule HuR inhibitor KH-3 as an experimental probe together with its inactive structural analog KH-3B as a control. *In vivo* experiments were performed in a HDM-induced airway inflammation mouse model, accompanied by *ex vivo* studies using murine splenic and lung CD4⁺ T cells, human lung CD4⁺ T cells, and human PBMC-derived ILC2s.

Single-cell transcriptomic data from bronchoalveolar lavage fluid (BALF) obtained from allergen-challenged asthmatic patients were reanalyzed to evaluate the expression of *ELAVL1* and *GATA3* genes across type 2 inflammatory cell populations in human asthma.

The work comprised four integrated experimental components: (1) HDM mouse model, (2) human lung CD4⁺ T cells, (3) human PBMC-derived ILC2, and (4) BALF scRNA-seq data from asthmatics. All mouse experiments used littermate controls with block randomization by sex and body weight (±2 g). Experimenters were blinded to treatment. No statistical methods were used to predetermine sample size for human studies, which were designed as mechanistic experiments based on prior HuR work. No outliers were excluded; all data are shown. Data collection and analysis of human subjects were not performed in a completely randomized manner due to biological constraints such as donor availability.

### Sex as a biological variable

Our prior studies including both male and female mice did not reveal significant sex-specific differences in the measured outcomes in established murine models of allergic airway inflammation; therefore, in the current study, in vivo experiments were conducted primarily in female mice. These findings are interpreted with the understanding that key pathways underlying type 2 inflammation are largely conserved across sexes.

For human studies, samples were obtained from both male and female donors when available; however, sex was not used as a stratification variable because of sample size limitations and the mechanistic nature of the study.

### Human subjects and ethics

Human donor selection for the PBMC studies followed the framework used in our prior HuR asthma study, with subjects classified by asthma status and type 2 inflammatory endotype when applicable (26). In the earlier work, type 2-high asthma was defined by blood eosinophilia and/or elevated fractional exhaled nitric oxide, and subjects receiving biologic therapy were excluded (26). For the present study, healthy control and asthmatic donor samples were processed using matched experimental workflows to minimize technical variation.

Human lung tissue used for lung CD4⁺ T cell studies was obtained from Gift of Life at the University of Michigan under IRB-exempt de-identified tissue protocols. PBMC specimens used for ILC2 studies were obtained from participants enrolled under an approved VA Merit study (CX002491, PI: Atasoy; IRBNet ID: ACORP 1644326), and written informed consent was obtained from all living participants in accordance with 45 CFR 46.116.

Asthma inclusion criteria for PBMC donor selection followed GINA 2025 type 2-high definitions, including blood eosinophils ≥300/μL and/or FeNO ≥25 ppb, no biologic therapy for ≥6 months, and FEV1/FVC <0.7. Control donors were age- (±5 years) and sex-matched, with FEV1 >80% predicted. Additional inclusion and exclusion criteria for type 2-high asthmatic and control subjects are provided in Supplementary Table 1.

For single-cell RNA-seq analyses, BALF scRNA-seq datasets from allergen-challenged asthmatic subjects were re-analyzed as described here. Human subject oversight, IRB approval, and informed consent procedures for these datasets were reported previously by Siddiqui *et al.* (47).

### Animals and housing

Six- to eight-week-old C57BL/6J mice of both sexes (The Jackson Laboratory; strain #000664) were used. Mice were housed in the University of Michigan Unit for Laboratory Animal Medicine specific-pathogen-free facility under standard laboratory conditions (12-hour light/dark cycle, controlled temperature and humidity) in individually ventilated cages with free access to standard chow and automated LIXIT-supplied drinking water. All procedures were performed in accordance with the U.S. National Institutes of Health Guide for the Care and Use of Laboratory Animals and were approved by the University of Michigan ULAM Institutional Animal Care and Use Committee (IACUC protocol PRO00011676).

### HDM-induced allergic airway inflammation model

Allergic airway inflammation was induced using a standardized house dust mite (HDM) extract from Dermatophagoides pteronyssinus (XPB82D3A2.5; Stallergenes Greer, Lenoir, NC, USA; lot 431619). According to the manufacturer, this lyophilized whole-body extract is prepared by bi-level extraction (1:20 and 1:10 w/v in 0.01 M ammonium bicarbonate, dialyzed against distilled water) and supplied as a lyophilized cake containing 20 mg dry weight, 6.05 mg protein, and 55.65 µg Der p 1 per 2.5-mL vial, with an endotoxin content of 20,875 EU/vial. Lyophilized HDM was reconstituted in sterile PBS to a stock concentration of 2.5 mg/mL, aliquoted, and stored at −20 °C until use. Mice were sensitized intraperitoneally on days 0 and 7 with 200 μg HDM in 100 μL sterile PBS; control mice received PBS alone. For allergen challenge, mice were administered 50 μg HDM in 50 μL PBS via the oropharyngeal (intratracheal) route on days 14 and 16 (63). KH-3 (HY-134601, MedChemExpress) was administered during the challenge phase at 100 mg/kg intraperitoneally, a dose selected based on tolerability and previously published safety data. KH-3 was prepared by initial dissolution in 5% ethanol and 5% Tween-80, followed by dilution in PBS. Mice received KH-3 or vehicle every other day from day 8 to day 16. Animals were euthanized 24 hours after the final HDM challenge for collection of BALF and lung tissue or were assessed for airway function as described below. Lungs designated for histology and imaging were collected following transcardial perfusion with 10% neutral buffered formalin (Epredia).

### Bronchoalveolar lavage and cell counts

Mouse lungs were lavaged with 1 mL PBS containing 0.5 mM EDTA. BALF was pooled and centrifuged at 300 g for 5 min at 4°C. Supernatants were aliquoted and stored at −80°C for cytokine analyses. Cell pellets were resuspended and counted using a Neubauer hemocytometer.

### Lung histology and inflammatory scoring

The left lung or designated lobes were fixed in 10% neutral-buffered formalin, paraffin-embedded, and sectioned at 5 μm. Sections were stained with hematoxylin and eosin to assess peribronchial and perivascular inflammation. Representative images were captured using a Leica DM IRB microscope equipped with an Olympus DP70 digital camera at 20× magnification.

### Cytokine measurement

BALF supernatants, lung homogenates, and activated splenic CD4⁺ T-cell culture supernatants from mice were analyzed using DuoSet ELISA kits (R&D Systems) for IL-4, IL-5, IL-13, and IL-17 according to the manufacturer’s instructions. Cytokine concentrations in lung homogenates were normalized to total protein when appropriate. For human cell culture supernatants, cytokines were quantified using the Human Th9/Th17/Th22 Luminex Performance Assay 18-plex fixed panel (LKTM009B, R&D Systems) according to the manufacturer’s instructions. This magnetic bead–based multiplex assay detects IL-9, CCL20/MIP-3α, CD40 ligand/TNFSF5, GM-CSF, IFN-γ, IL-1β, IL-2, IL-4, IL-5, IL-6, IL-10, IL-12p70, IL-13, IL-15, IL-17A, IL-17E/IL-25, IL-33, and TNF-α.

### Airway physiology

Airway hyperresponsiveness was assessed 24 hours after the final HDM challenge in anesthetized, tracheostomized mice by plethysmography using a Buxco FinePointe resistance/compliance system (Data Sciences International, Wilmington, NC). Total respiratory system resistance (Rn) was recorded in response to increasing doses of nebulized methacholine (0, 10, 20, and 40 mg/mL) delivered via the ventilator, and resistance values (Rn) were analyzed across the methacholine dose–response curve, as described earlier (64).

### Isolation and stimulation of murine CD4⁺ T cells

Splenic CD4⁺ T cells were isolated by magnetic bead separation (Miltenyi Biotec). MACS-purified CD4⁺ T cells were cultured in complete media consisting of RPMI 1640 (Gibco) supplemented with 10% heat-inactivated FBS, 2 mM L-glutamine, 1 mM sodium pyruvate, 55 μM β-mercaptoethanol, and gentamicin. Cells were pretreated with KH-3 or inactive analogue KH-3B (2 μM, for 2 hours) and then stimulated with plate-bound anti-CD3 plus soluble anti-CD28 (both from Invitrogen) for up to 4 days. For short-term assays, cells were harvested after 4 days of stimulation for intracellular cytokine staining by flow cytometry and for measurement of secreted cytokines in culture supernatants by ELISA (26).

### Human lung CD4⁺ T-cell isolation and activation

Fresh de-identified human lung tissue (University of Michigan Lung Biorepository or Gift of Life) was mechanically and enzymatically dissociated to obtain single-cell suspensions, digested in collagenase A and DNase I (both from Roche Diagnostics) at 37°C, processed on a gentleMACS Dissociator, and filtered through 70 μm strainers. CD4⁺ T cells were enriched using Human CD4 MicroBeads on LS columns (Miltenyi) in MACS buffer, achieving >95% purity by CD3/CD4 flow cytometry. Purified CD4⁺ T cells were cultured in complete media, pretreated for 2 hours with KH-3 or KH-3B, and then stimulated for 4 days on plates coated with anti-CD3 (5 μg/mL) and anti-CD28 (2 μg/mL). Supernatants were collected for multiplex cytokine assays, and cell pellets were used for RNA extraction and qPCR; in selected experiments, actinomycin D; ActD (5 μg/mL) was added after 4 days of activation to assess mRNA decay.

### Human PBMC-derived ILC2 isolation, culture, and stimulation

Cryopreserved PBMCs from healthy controls and type 2–high asthmatic donors were thawed and recovered in parallel under identical conditions. PBMCs were originally isolated from EDTA-anticoagulated blood by density-gradient centrifugation over Ficoll-Paque PLUS (17-1440-03, GE Healthcare; 16 mL blood layered over 12 mL Ficoll, centrifuged at 2000 ×g for 40 min at room temperature with no brake). After thawing, PBMCs were rested overnight in complete medium supplemented with 10 ng/mL IL-7.

ILC2s were isolated using the Human ILC2 Isolation Kit (Miltenyi Biotec, 130-114-825) according to the manufacturer’s instructions. Peripheral blood mononuclear cells were first incubated with a biotin-conjugated lineage antibody cocktail containing antibodies against CD2, CD3, CD14, CD16, CD19, CD56, CD235a (Glycophorin A), and CD123, followed by Anti-Biotin MicroBeads, and lineage-positive cells were depleted over LS MACS columns. The resulting lineage-negative fraction was then stained with CD294 (CRTH2)-PE and labeled with Anti-PE MicroBeads, and CD294⁺ cells were positively selected on MS MACS columns to enrich for ILC2s. This two-step procedure routinely yielded lineage-negative, CD294⁺ cells at approximately 3 × 10³ cells per 50 mL blood. For flow cytometric characterization, enriched cells were further confirmed to be Lin⁻CD45⁺CD127⁺CD161⁺CD294⁺.

Purified ILC2s were cultured in complete medium, pretreated with KH-3 or the inactive analog KH-3B for 2 hours, and then stimulated with recombinant human IL-7, IL-33, and TSLP (10 ng/mL each) for 4 days without compound washout. Cell-free supernatants were collected and stored at −80 °C for multiplex cytokine (Luminex) analysis, and cells were harvested for RNA extraction and qPCR.

On day 4, parallel cultures were subjected to ActD mRNA decay assays. At time 0 (immediately before ActD addition), supernatants were collected for Luminex analysis. Cultures were then treated with ActD and incubated for 4 hours, followed by collection of supernatants. Cells were lysed directly in RLT buffer and stored at −80 °C until RNA isolation and downstream qPCR analysis.

### RNA extraction, qPCR, primer sequences, and mRNA stability

Total RNA from murine and human cells (including CD4⁺ T cells and ILC2s) was isolated using RNeasy Mini kits (Qiagen) according to the manufacturer’s instructions. RNA integrity was verified by spectrophotometric analysis (NanoDrop), and only samples with acceptable quality metrics (A260/A280 ratios of 1.8–2.1 and A260/A230 ratios ≥1.8) were used for downstream analyses.

For standard expression analysis, cDNA was synthesized using the High-Capacity cDNA Reverse Transcription Kit (Applied Biosystems, catalog 4368814) with 10 ng–1 μg RNA per reaction, depending on sample type. Quantitative PCR was performed on a QuantStudio 3 Real-Time PCR System (Applied Biosystems, Foster City, CA) using SYBR Green chemistry with validated primer pairs (Integrated DNA Technologies; HPLC-purified, 500 nM final primer concentration). Relative mRNA expression was calculated using the ΔΔCt method, with *Gapdh* (mouse) or *GAPDH* (human) serving as the internal reference gene.

For mRNA stability assays, mouse and human cells were treated with ActD (5 μg/mL) to block transcription after the indicated activation period, following our previously published protocol (49). RNA was harvested at serial time points after ActD addition (0-4 hours), converted to cDNA, and target transcript abundance was quantified by qPCR as described above. For each transcript, values at time zero were set to 100%, and remaining mRNA at subsequent time points was expressed relative to this baseline; decay curves were fitted using nonlinear regression, and mRNA half-life (t₁/₂) was calculated from the best-fit model.

Primer sequences and amplicon sizes for all mouse and human targets are listed in Tables 1 and 2, respectively.

**Table 1.**
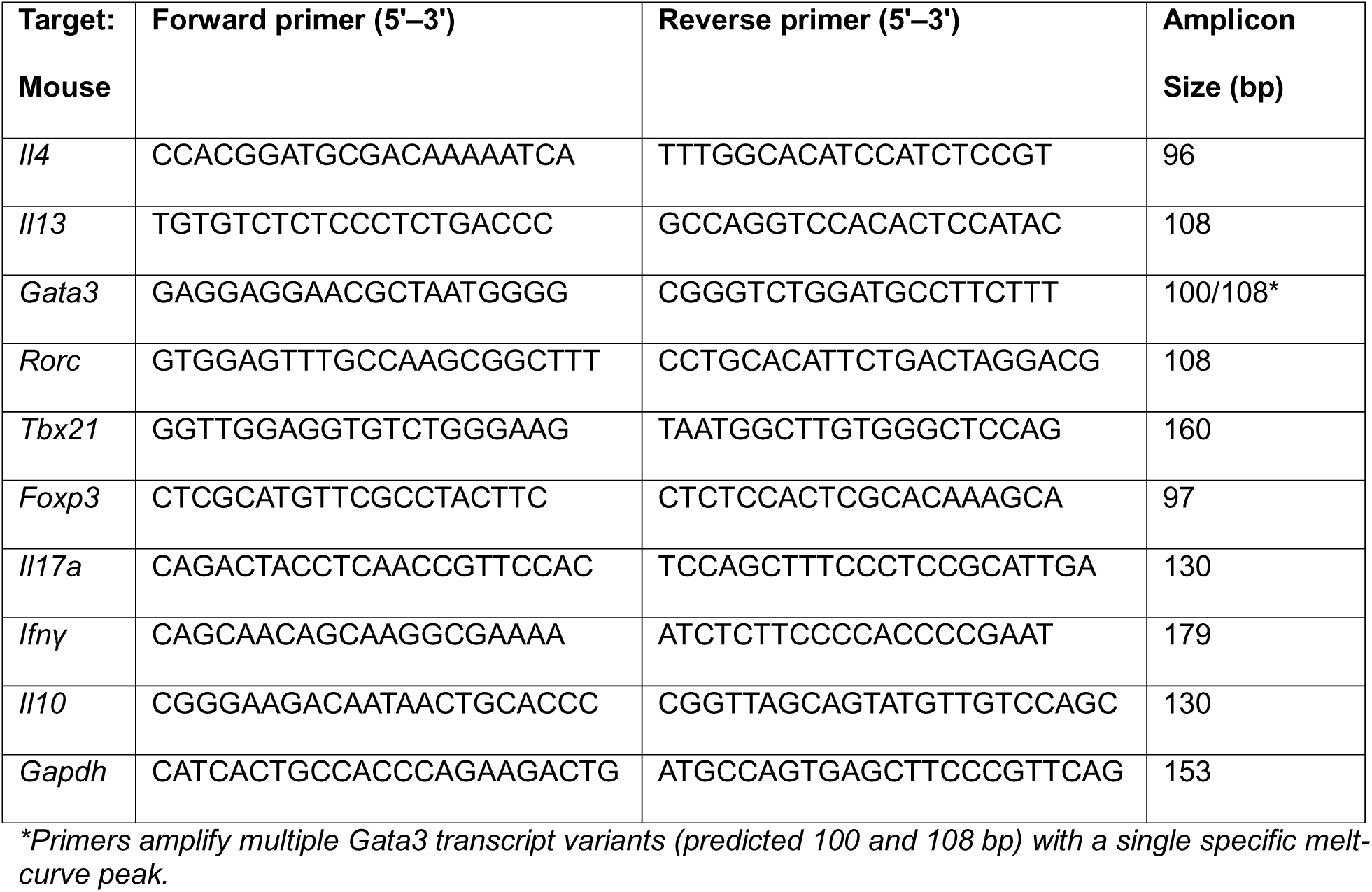
Mouse qPCR primer sequences and amplicon sizes.

**Table 2.**
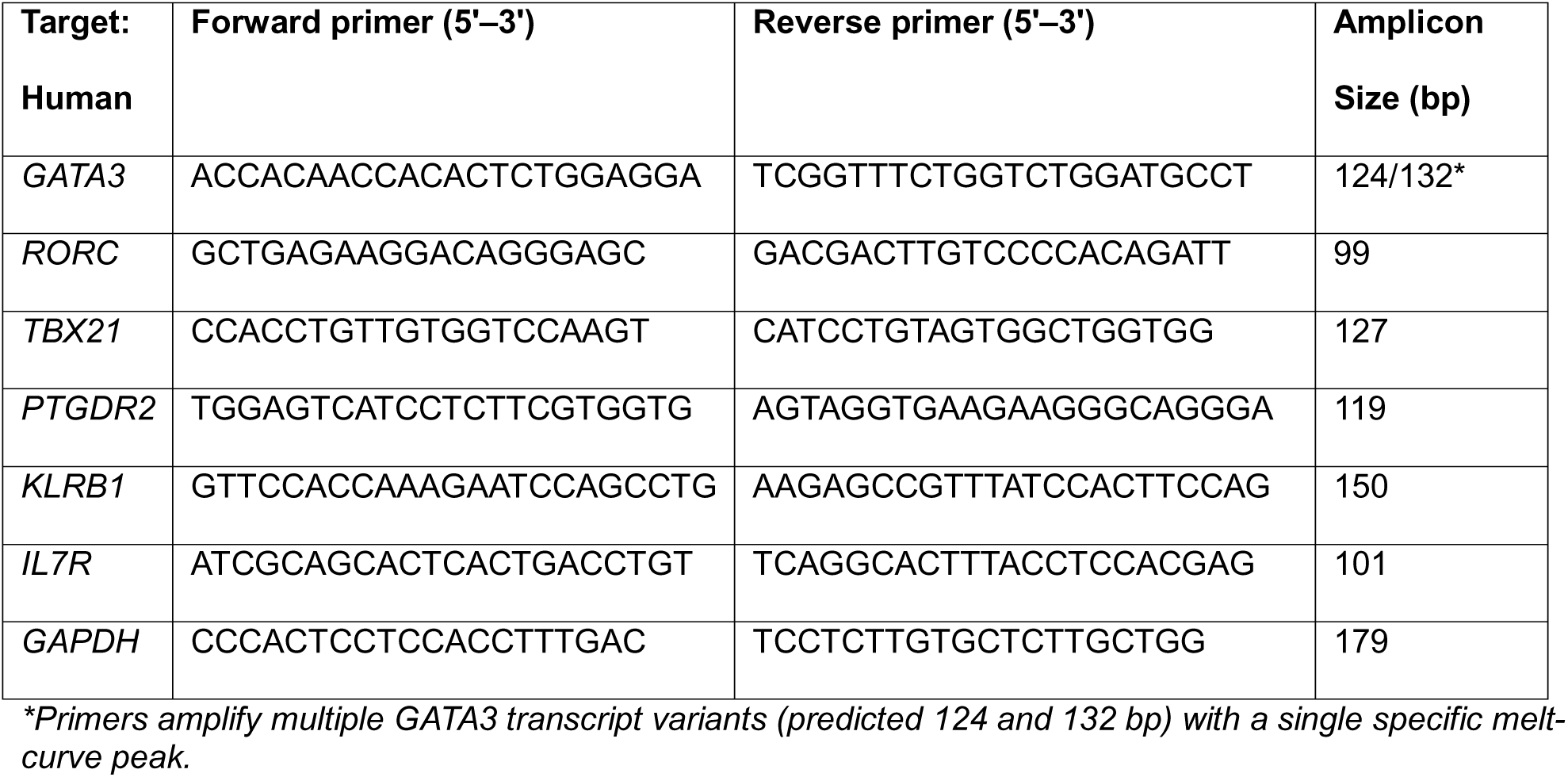
Human qPCR primer sequences and amplicon sizes.

### Flow cytometry

CD4⁺ T cells were isolated from spleens of C57BL/6 mice and activated *in vitro* for 4 days with plate-bound anti-CD3 and soluble anti-CD28 before processing for flow cytometry. Prior to staining, cells were incubated with TruStain FcX Fc receptor–blocking solution (BioLegend) to minimize nonspecific binding and then labeled with LIVE/DEAD™ Fixable Dead Cell Stain (Invitrogen) compatible with the multicolor panel. Surface staining was performed at 4°C for 30 minutes in the dark using fluorochrome-conjugated antibodies against CD3 and CD4. For intracellular staining of cytokines and GATA3, cells were fixed and permeabilized using an intracellular staining buffer set (eBioscience), followed by staining with fluorochrome-conjugated anti-cytokine and anti-GATA3 antibodies. Fluorescence-minus-one, single-color, and isotype controls were included for gate setting and compensation, and UltraComp eBeads (Invitrogen) were used for compensation. Data were acquired on a BD FACSCanto flow cytometer and analyzed using FlowJo software (Tree Star) (49).

### Single-cell RNA sequencing

BALF single-cell RNA-seq datasets generated after segmental allergen challenge were obtained from Siddiqui *et al*. (47) and reanalyzed in the present study. Briefly, data generated using the 10x Genomics Chromium platform and Illumina sequencing were analyzed in Seurat, including normalization, dimensionality reduction, clustering, and differential expression testing with false-discovery-rate correction. *ELAVL1* (encoding HuR) and GATA3 expression were examined across immune cell clusters to assess enrichment in Th2-associated populations.

### Statistical analysis

Unless otherwise specified, data are presented as mean ± SD, as indicated in figure legends. Comparisons between two groups were performed using unpaired or paired Student’s t tests. Multiple-group comparisons were analyzed by one-way or two-way ANOVA with appropriate post hoc tests, as indicated in figure legends. For scRNA-seq, differential expression was assessed using Wilcoxon rank-sum tests with FDR correction (FDR < 0.05) (47). Exact p values are reported when feasible; statistical significance was defined as *p <* 0.05.

## Supporting information

Graphical Abstract

## Data availability

All data generated or analyzed during this study are included in this article and its supplementary information files. Raw datasets are available from the corresponding author upon request.

## Author contributions

F.F. designed the research studies, conducted experiments, acquired and analyzed data, and wrote the manuscript. L.Y., J.H., B.T., K.H., J.M., and K.B. conducted experiments and acquired and analyzed data. S.N. acquired and analyzed data. L.X. contributed to study design and provided reagents. M.H. and S.H. contributed to data interpretation and provided reagents and tools. U.A. conceived and supervised the study, contributed to study design, provided funding and reagents, and contributed to manuscript writing.

## Acknowledgments

The authors thank Fahimeh Fattahi for guidance with qPCR optimization and primer design, Natalie Walker for assistance with intratracheal administration in the HDM mouse model, and Julia Khater for advice on human lung tissue processing and handling.

