## Supplementary figures and images for "HuR Regulates GATA3-Driven Type 2 Inflammation in CD4^⁺^ T cells and ILC2 in Airway Inflammation"

### Graphical Abstract

#### Graphical Abstract

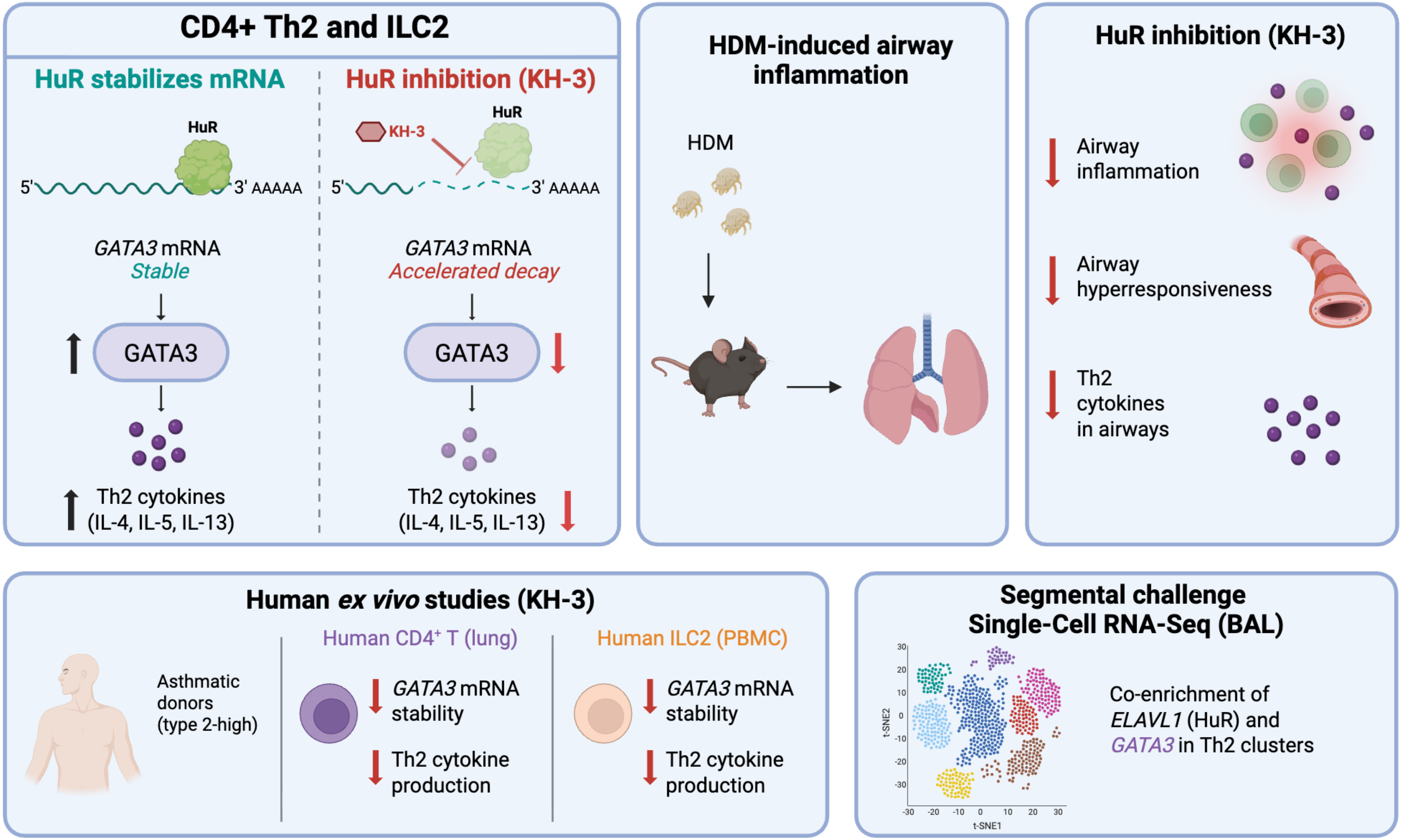
