## Supplementary Table 1 for "HuR Regulates GATA3-Driven Type 2 Inflammation in CD4^⁺^ T cells and ILC2 in Airway Inflammation"

**Supplementary Table 1. Criteria for Recruiting Type 2 Asthmatic and Control Individuals for the Study**

| **Group** | **Inclusion Criteria** | **Exclusion Criteria** |
| --- | --- | --- |
| **Type 2–high Asthma** | • Male or female Veterans • Age 18–75 years • No other chronic lung disease • Diagnosed Type 2–high asthma, defined by ≥1 of the following:    – Eosinophils ≥300/µL    – FeNO ≥25 ppb    – IgE >30 IU/mL    – Sensitization to ≥2 aeroallergens (skin prick or RAST) • No biologic therapy within the past 6 months | • COPD or bronchiectasis • Other chronic lung disease • BMI >35 • Pregnancy • Recent infection (within 6 weeks) • Unstable cardiac disease • Severe OSA with hypoventilation • Contraindication to anticoagulation • FEV₁ <60% predicted • Active tuberculosis • History of fibrotic lung disease • Injection drug use or prior thoracic radiation • Regular cannabis or e-cigarette use • PREPARE-triggering comorbidities without clearance |
| **Healthy Controls** | • Male or female Veterans • Age 18–75 years • No chronic lung disease • Non-atopic • Normal pre-bronchodilator spirometry:    – FEV₁ ≥80% predicted    – FEV₁/FVC > LLN • Eosinophils <150/µL • FeNO <25 ppb • BMI <35 • No tobacco use (lifetime <100 cigarettes; ≥6-month abstinence from tobacco/vaping) | • Same exclusion criteria as asthma group • Any history of asthma or allergic disease |
